# A weak Foxp3 hypomorph enhances spontaneous and therapeutic immune surveillance of cancer in mice

**DOI:** 10.1101/570671

**Authors:** José Almeida-Santos, Marie-Louise Bergman, Inês Amendoeira Cabral, Vasco Correia, Íris Caramalho, Jocelyne Demengeot

**Affiliations:** Instituto Gulbenkian de Ciência

## Abstract

It is well established that therapeutic impairment of Foxp3^+^ regulatory T cells (Treg) in mice and humans favors immune rejection of solid tumors. Less explored are the genetic associations between Foxp3 allelic variants and tumor incidence, only sporadically reported in human studies. In this work, we tested and demonstrate that Foxp3^fGFP^, an allele classified as hypomorphic in Th1 inflammatory contexts but not affecting health at steady state, confers increased anti-tumor immunity. Our conclusions stem out of the analysis of three tumor models of different tissue origin, in two murine genetic backgrounds. When compared to wild type animals, mice carrying the Foxp3^fGFP^ allele spontaneously delay, reduce or prevent primary tumor growth, decrease metastasis growth and potentiate the response to anti-CTLA4 monotherapy. These findings suggest that allelic variance at the Foxp3 locus may have significant impact on cancer incidence and/or the success of cancer-immunotherapies in humans.

## Introduction

Regulatory T cells (Treg), a subset of CD4^+^ cells, express the transcription factor Foxp3 that defines a transcriptional profile essential for their differentiation and function (1). By controlling the activation of conventional T cells, Treg guarantee the establishment and maintenance of immune tolerance to self-components (2). It is also well established that depletion or inhibition of Treg in mice and humans, favors immune rejection of solid tumors (3, 4). Several allelic variants in humans have been associated with various autoimmune diseases (5) and loss of function mutations in the Foxp3 gene are responsible for the fatal IPEX syndrome(6). Foxp3 allelic variants were also associated with increased susceptibility to colorectal and non-small cell lung cancer, and progression of breast cancer (5). However, Foxp3 expression is not restricted to Treg and acts as a cell intrinsic tumor suppressor that represses the oncogenes SKP2, HER2 (7, 8) or cMYC (9) in solid tumors. Thus, it remains unclear whether allelic variants of the Foxp3 gene can affect immune surveillance of cancer. In turn, it is conceivable that protective Foxp3 alleles may also enhance the effectiveness of immune-therapies for cancer.

The development of Foxp3 reporters in mice fortuitously generated Foxp3 alleles that are functionally impaired, to various degrees (10–13). The commonly used Foxp3^fGFP^ knock-in allele, that encodes a Foxp3 protein fused at its N-terminus to the enhanced green fluorescence protein (eGFP), moderately alters the transcriptional signature and phenotype of Treg, with functional impacts in models of spontaneous autoimmunity and infection (11, 12, 14). As the Foxp3^fGFP^ allele does not affect health at steady state in the reference C57Bl/6 (B6) genetic background (12), it is an ideal model to test for specific effect on tumor progression. To dissociate tumorigenesis from anti-cancer immunity, we used three transplantable tumor models. We evidence reduced primary tumor growth associated with increased immune responses, reduced metastatic progression and enhanced response to anti-CTLA4 monotherapy in Foxp3-GFP mice when compared to wild-type (WT) controls. This preclinical analysis supports the notion that allelic variance at the Foxp3 locus may serve as predictive indicators for personalised therapy and prognostics, to the benefit of cancer patients.

## Results and Discussion

### The Foxp3^fGFP^ allele enhances spontaneous immune-surveillance of primary tumors

The Foxp3^fGFP^ allele on a B6 background has been reported to affect Treg phenotype and function but not health at steady state (12). We generated BALB/c (Ba) Foxp3-GFP mice which, compared to gender and age matched WT animals, have slightly underrepresented Treg that express increased Foxp3 protein (as reported for mice on the B6 background (11)), bear normal numbers of activated or interferon-γ (IFN-γ) producing T cells (Fig. S1) and do not show sign of disease.

To test whether the Foxp3^fGFP^ allele provides for enhanced anti-tumor immunity, we chose to monitor the CT26 colorectal carcinoma cell line, derived from a BALB/c mouse. While CT26 engraftment and growth is successful in WT animals, its intrinsic immunogenicity is readily revealed upon depletion of Treg by administration of diphtheria toxin (DT) in DEREG mice, either one (15) or two weeks (Fig. 1A) after implantation. In both cases, the tumors were consistently and fully rejected, confirming Treg restrain a potentially vigorous immune control of CT26 tumors, and validating the choice of this tumor model to test the impact of Foxp3 allelic variants in cancer immune-surveillance. Of note, in this experimental setting as in others, treatments of WT mice with DT is not innocuous (compare left panels Fig. 1A and 1B, and below Fig. 2A and 2C). We therefore monitored the growth of CT26 cells injected subcutaneously (s.c.) in Ba.Foxp3-GFP or WT mice (representative experiment in Fig. 1B). The experimental and control groups were readily and reproducibly distinguishable, with delayed and reduced tumor growth in the former. Due to the heterogeneous shapes of each tumor growth curve inside the experimental groups, pooled raw data were treated through the Tumor Control Index (16) that informs on tumor regression, stability and rejection, to infer more faithful statistical analysis (Fig. 1H). Strikingly, while all WT animals allowed engraftment and regular tumor size progression, CT26 growth was either delayed or fully prevented in more than a third of the Foxp3-GFP mice. To ascertain that immune responses were enhanced in Foxp3-GFP mice, tumor infiltrating lymphocytes (TIL) were analyzed 15 days post-implantation (Fig. 1C-E and S2). The tumor weight was reduced (Fig. 1C), the frequency of CD8 IFN-γ producing cells was increased (Fig. 1D) and the ratio Treg to CD8 was inverted (Fig.1E) in Foxp3-GFP compared to WT mice. In the tumor draining lymph node (DLN), where Treg were not overrepresented, these differences were not found in the draining lymph nodes (Fig. S2). Overall, these data demonstrate that the Foxp3^fGFP^ allele, a fusion of eGFP at the N-terminal part of Foxp3, potentiates spontaneous anti-tumor immunity without affecting health in the BALB/c reference strain. These results resonate with recent findings from the Mathis-Benoist’ lab (17). By introducing distributed alanine replacement mutations in the mouse Foxp3 gene, they identified two mutant alleles that conferred B6 mice with reduced growth of the MC38 primary tumor. One of these alleles does not affect health at steady state and is located in the N-terminal proline-rich region of Foxp3, where no specific motifs have been identified.

**Figure 1.**
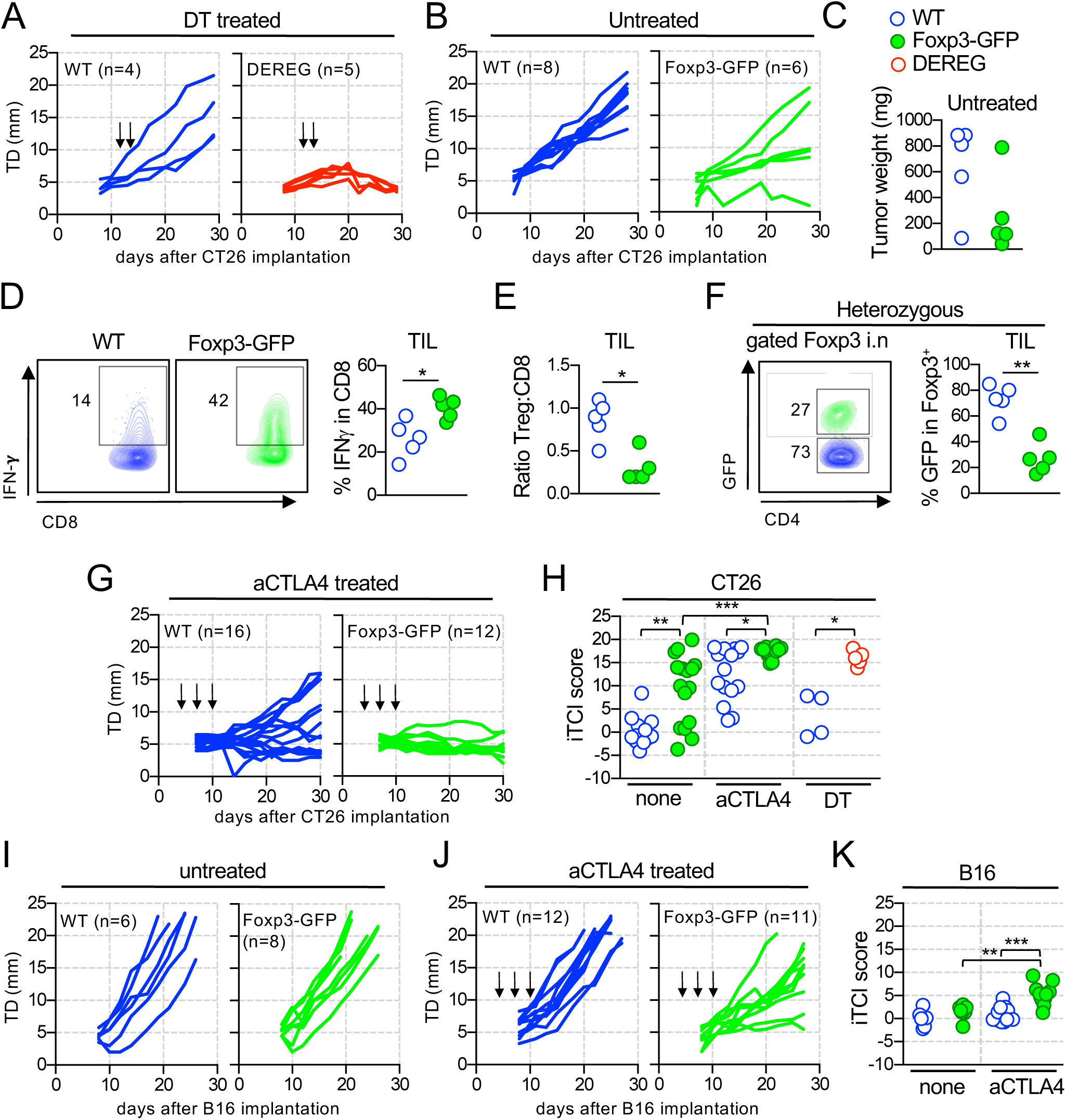
The Foxp3^fGFP^ allele promotes spontaneous and therapeutic tumor immune-surveillance. 3×10^5^ CT26 (A-H) or 2×10^5^ B16 cells (I-K) were injected s.c. in the flank of mice with a BALB/c or B6 genetic background, respectively. **A**) Tumor diameter (TD) along time in WT (left) and DEREG (right) littermates, treated with 1µg of diphtheria toxin (DT) at day 13 and 14 post-implantation (arrows), p=0.0196. **B**) as in (A) in untreated WT and Foxp3-GFP mice, p=0.0052. Shown is 1 representative of 3 independent experiments **C-E**) Analysis of untreated WT and Foxp3-GFP mice (n=5 for each group) 15 days post-implantation for tumor weight (C) and for tumor infiltrating lymphocytes (TIL) (D-E). Shown is the frequency of interferon-γ (IFN-γ) producing CD8 cells (D) and the ratio of i.n. Foxp3^+^ (Treg) to CD8^+^ cells (E). **F**) Analysis of the TIL in heterozygous Foxp3^fGFP/wt^ mice 15 days post-implantation. Shown is the frequency of GFP expressing cells in gated i.n. Foxp3^+^ cells. **G**) as in (B), except that mice received 100µg of aCTLA4 at day 4, 7 and 10 post tumor implantation. Shown are 2 independent experiments, pooled, p=0.0449. **H**) Individual Tumor Control Index (iTCI) scores for CT26 growth curves presented in (A, B and G), pooling independent experiments for each group. **I, J**) As in (B) and (G) for B16 tumors and mice on a B6 background. For (J), shown are 2 independent experiments, pooled, p=0.0041. **K**) iTCI scores for B16 growth curves shown in (I) and (J). All statistical analysis shown inside the panels were performed using nonparametric Mann-Whitney test: *P < 0.05, **P 0.005 and ***P < 0.001. Statistics for tumor growth (p value inserted in the legend) used two-way ANOVA.

**Figure 2.**
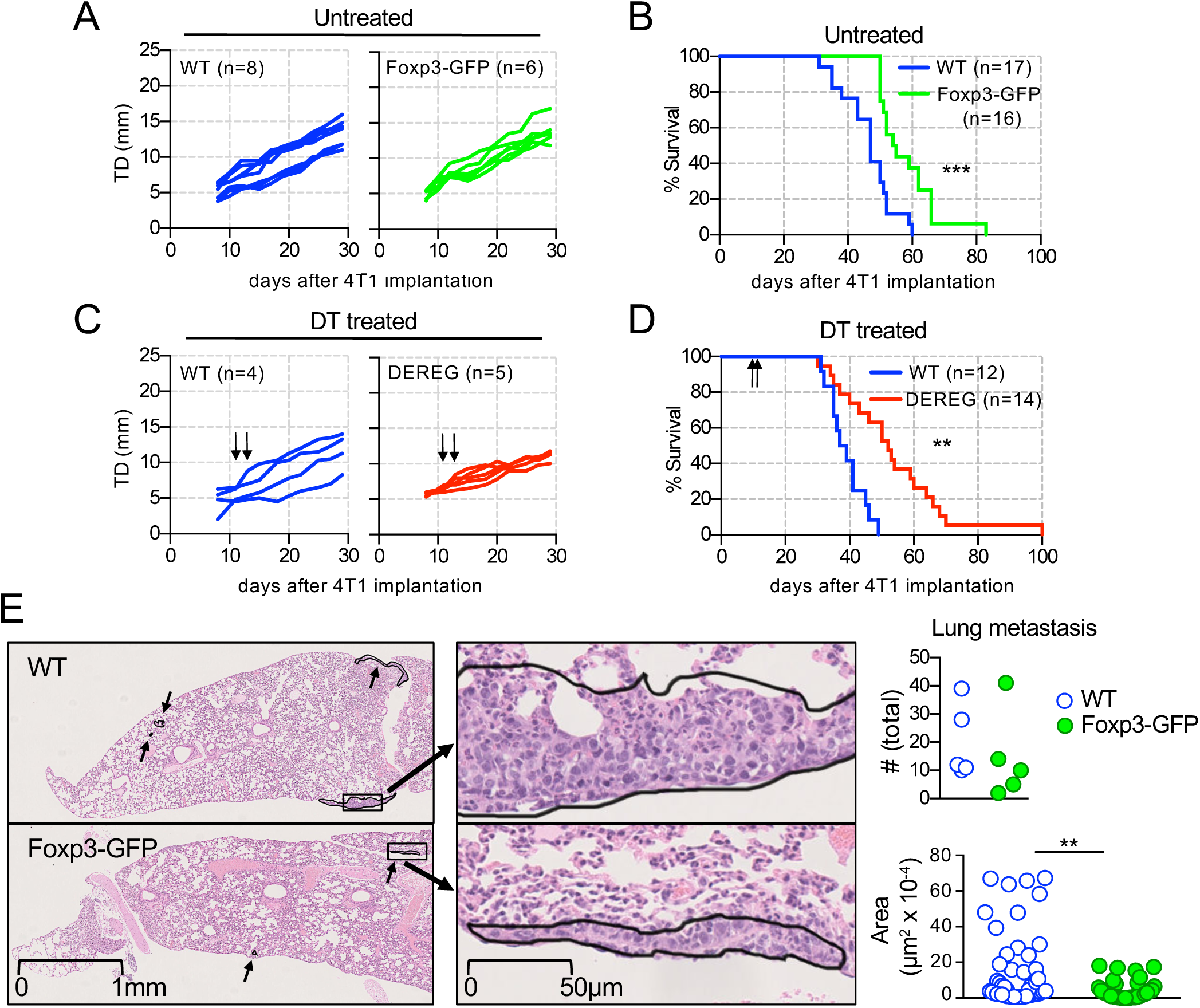
The Foxp3^fGFP^ allele reduces metastasis progression. Mice on a BALB/c background were injected s.c. with 5×10^4^ 4T1 cells. **A, B**) Tumor growth (A) and mice survival (B) in WT and Foxp3-GFP mice. **C-D**) Tumor growth (C) and survival in WT and DEREG littermates treated with DT as in Fig1. **E**) Left, representative histological analysis of the lungs from WT and Foxp3-GFP mice 24 days post-implantation, in lower magnification pictures the thin arrows point at metastatic foci, the thick arrows indicate the boxed areas that are presented also at high magnification. Right, metastasis quantification and area estimates. For the latter, plotted is the individual area of each metastatic focus identified in 5 mice, for each genotype. Statistical analysis was performed using two-way ANOVA for tumor growth (ns for both A and C), logrank test for survival curves, and Nonparametric Mann-Whitney test for metastasis number and area.

We next tested whether the Foxp3^fGFP^ allele confers a disadvantage to Treg in the CT26 context. Owing to the fact that Foxp3 is located on the X chromosome, heterozygous Foxp3^fGFP/WT^ females are mosaic and mimic a cell competition assay. Analysis of TIL 15 days post-implantation of CT26 into Foxp3^fGFP/WT^ females revealed that a large majority of Treg expressed the WT allele (Fig. 1F), indicating that Treg expressing the Foxp3^fGFP^ allele are less fit. A similar trend was found when analyzing the dLN (Fig. S2). Although Bettini et al did not address the impact of the Foxp3^fGFP^ allele on tumor growth, they reported a similar disadvantage for Treg expressing the Foxp3^fGFP^ allele when analyzing B6.Foxp3^fGFP/WT^ females implanted with the B16 melanoma (12). Cancer immunosuppression is a complex process, for which both thymic derived (tTreg) (18) and peripherally differentiated Treg (pTreg) (19) have been incriminated, and both cell subsets were shown to be affected by the Foxp3^fGFP^ allele (12, 13).

### The Foxp3^fGFP^ allele enhances the effectiveness of cancer immunotherapies

We next tested whether the Foxp3^fGFP^ allele potentiates therapeutic response to anti-CTLA4 mAb (aCTLA4) treatment. We administered aCTLA4 i.p. during the first week following tumor implantation in WT and Foxp3-GFP mice, and followed subsequent tumor progression (Fig. 1G-K). As previously reported (20), WT mice implanted with CT26 partially responded to aCTLA4 monotherapy, with only a fraction of them rejecting or delaying tumor growth. In contrast, all treated Foxp3-GFP mice prevented tumor growth, with most animals maintaining a tumor free state 1-month post-implantation (Fig. 1G). These dramatic results encouraged us to test B6.Foxp3-GFP mice implanted with the poorly immunogenic B16 melanoma for which aCTLA4 monotherapy has been reported ineffective (20). Strikingly, although the Foxp3^fGFP^ allele by itself did not affect B16 engraftment or progression (Fig. 1I), aCTLA4 treatment delayed tumor growth in the majority of Foxp3-GFP mice but not in WT controls (Fig. 1J, K). Finally, enhanced therapeutic response afforded by the Foxp3^fGFP^ allele was not accompanied by overt systemic effects as indicated by constant body weight (Fig. S2).

Our finding that the Foxp3^fGFP^ allele potentiates aCTLA4 therapy, together with the evidence that aCTLA4 kills Treg (21, 22), echoes with a previous report indicating that the same allele amplifies the effectiveness of DT treatment in DEREG mice in an infection setting (14). Moreover, the improved efficacy of aCTLA4 treatment that we evidence in Foxp3-GFP mice, both in BALB/c and in B6 backgrounds, suggests that allelic variance at the Foxp3 or other Treg signature genes may discriminate responder from non-responder patients submitted to this therapy.

### The Foxp3^fGFP^ allele delays metastasis dissemination

The last phase of tumor progression is metastatic dissemination, a process that is well modelled by the 4T1 breast carcinoma derived from a BALB/c mouse. The 4T1 tumor grows slower than CT26 or B16 at the site of implantation, resists many immune interventions and is highly metastatic with an average mean survival of the host of 50 days post-implantation (23). A moderate role for Treg in facilitating 4T1 primary tumor growth (15) and precipitating death (24) has been reported. Monitoring WT and Foxp3-GFP animals implanted s.c. with 4T1 cells revealed similar growth of the primary tumor while survival was significantly prolonged in the latter group (Fig. 2A, B and S3). Prolonged survival was also observed in DEREG animals administrated with DT, as mean to induce a transient depletion of Treg, early after 4T1 implantation (Fig 2C, D and S3). We next ascertained that the prolonged survival in Foxp3-GFP mice bearing 4T1 tumors associated with reduced metastasis dissemination. We first confirmed that resection of the primary tumor during the second week post-implantation, but not later, greatly enhances survival (Fig. S3), an intervention shown by others to prevent the metastatic process (23). This result suggested that analysis at 3 weeks post implantation would be suitable to quantify metastasis dissemination in WT and Foxp3-GFP mice. Lungs were harvested and a histological assessment of metastasis number and size was performed (Fig. 2E). Although metastatic foci were similar in number in both groups of mice, large nodules were only found in WT animals. Together, these findings indicate that the Foxp3^fGFP^ allele restrains the dissemination stage of 4T1 tumors and suggest that identification of allelic variants at the Foxp3 or downstream genes may serve to guide prognostic estimates for cancer patients.

## Concluding remarks

Of relevance for experimental biology at large, our work provides further evidence that experimental variations can be related to reporter alleles, now in the context of cancer immune-surveillance. In the frame of immune tolerance, the evidence that Treg are essential components of immune regulation produced the notion that affecting Treg function for therapeutic reasons, such as for unleashing tumor immunity, will also unleash pathologic autoimmune reactivities. The adverse events tightly associated with tumor immunotherapies, and most notably with aCTLA4 therapy, are indeed autoimmune and inflammatory manifestations, often jeopardizing treatment continuation. In apparent contradiction with this notion, the Foxp3^fGFP^ allele on a B6 or BALB/c genetic background, does not compromise health at steady state and yet favors anti-tumor immunity. However, when introduced on a genetic background carrying several susceptibility alleles that together promote autoimmune Type 1 Diabetes, the Foxp3^fGFP^ allele precipitates disease (11, 12). While steady state mice are unlikely to be faithful models of humans continuously exposed to inflammatory triggers, it is conceivable that in the range of variations allowing for return to homeostasis, weak Foxp3 hypomorphs on an otherwise healthy genetic background would be beneficial to fight chronic infections and cancer, without tilting the balance toward autoimmunity. It therefore remains possible that weak but beneficial Foxp3 alleles are present in human population, more easily identifiable in males, and that genetic analysis would provide biomarkers to guide cancer prognostics and therapeutic strategies.

## Materials and Methods

### Mice

C57BL/6 Foxp3^tm2Ayr^ (B6.Foxp3-GFP, carrying the Foxp3^fGFP^ allele) were originally provided by A. Rudensky and backcrossed for at least 10 generations to the BALB/c.ByJ background (Ba.Foxp3-GFP). Foxp3-GFP mice in experiments are either homozygotes females or hemizygote males, unless otherwise specified. BALB/c-Tg(Foxp3-DTR/EGFP)23.2Spar (DEREG) were originally provided by T. Sparwasser and maintained as heterozygotes. C57BL/6J and BALB/c.ByJ served as WT controls. All mice were bred and raised at the Instituto Gulbenkian de Ciência (IGC) animal facility under specific pathogen–free conditions (SPF). Mouse experiments were conducted according to the Federation for Laboratory Animal Science Association guidelines and were approved by the ethic committee of the IGC.

### Cell lines and tumor models

Tumor cells CT26 and 4T1 (both ATCC) as well as B16-F10-luc2 (B16) (CaliperLS) were cultured at 37°C in RPMI 1640 (Life Technologies) supplemented with 10% Fetal Bovine Serum (Biowest), 100 U/ml Penicillin, 100 µg/ml Streptomycin, 50 µg/ml Gentamicin and 50 µM 2-Mercaptoethanol (all Life Technologies). After trypsin treatment, single cell suspensions were maintained in ice cold calcium- and magnesium-free HBSS (both Life Technologies). Mice were implanted with 3×10^5^ CT26, 2×10^5^ B16 or 5×10^4^ 4T1 cells through a subcutaneous injection of 100µl in the right flank. Tumor size was mesured with a caliper every 2 or 3 days, from day 8 post injection, and the tumor diameter (TD) calculated as TD=(L+W)/2. For ethical reasons, mice were sacrificed when TD≥20 mm. By the end of each experiment, rejection in animals scored as TD ≤5mm was confirmed upon dissection.

### Diphtheria toxin (DT) and anti-CTLA4 treatments

For Treg depletion, mice received 1 µg DT (322326-1, Calbiochem) diluted in 100 µl PBS through intraperitoneal (i.p.) injection, 13 and 14 days after tumor implantation. For anti-CTLA4 treatments, mice received 100 µL of PBS containing 100 µg of mAb 4F10, injected i.p., on days 4 and 7 and 10 post tumor implantation.

### Antibodies and FACS analysis

Single cell suspensions from mouse inguinal lymph nodes (iLN) were prepared by mechanical disruption in PBS (Life Technologies) containing 2% Fetal Bovine Serum (Biowest). Tumor infiltrating lymphocytes (TIL) were recovered after tumor digestion for 30 minutes in HBSS medium (Life Technologies) containing 10mM EDTA, 0,1% BSA, 1mg/ml collagenase type IV and 100µg/ml DNAse I, followed by separation on a Percoll gradient (all from Sigma). Live cells were counted using 10 µm latex beads (Beckman Coulter) and Propidium Iodide (Sigma). For cytokine analysis, cells were incubated for 4h at 37°C in RPMI 1640 (Life Technologies) containing phorbol 12-myristate-13-acetate and Ionomycin (both Sigma), and Brefeldin A (eBioscience). For intranuclear and intracytoplasmic staining, cells were first pre-incubated with Fc-block, stained for surface markers and then incubated overnight at 4°C in Foxp3 fix/permeabilization buffer (eBioscience). mAb are listed in Table 1. Samples were processed on a Cyan ADP instrument and analyzed with the FlowJo software.

**Table 1.** monoclonal Antibodies used in this study

| Reactivity | Clone | Conjugate | Supplier* |
| --- | --- | --- | --- |
| CTLA4 | 4F10 |  | In house |
| Fc (Fc-block) | 2.4G2 |  | In house |
| CD4 | GK1.5 | PB | In house |
| CD44 | IM7 | FITC | BD Pharmingen |
| CD44 | IM7 | e450 | eBioscience |
| CD62L | H1.2F3 | Biot/PE | BioLegend |
| CD8 | YTS169.4 | Biot/PB | In house |
| Foxp3 | FJK-16s | FITC/e450/APC | eBioscience |
| INF-gama | XMG1.2 | PE | BD Pharmingen |
| TCR-beta | H57-597 | PerCP-Cy5.5 | eBioscience |
| TCR-beta | H57-597 | A647 | In house |
| Streptavidin |  | APC-Cy7 | BioLegend |
| APC, allophycocyanin; Biot, biotin; Cy, cychrome; PB, Pacific Blue; PE, phycoerythrin; FITC, fluorescein isothiocyanate; e, eFluor; PerCP, Peridinin-Chlorophyll-protein |  |  |  |
| *All in house produced antibodies were purified on protein G columns. |  |  |  |

### Histological assessment

Tumor bearing mice were perfused with PBS, lungs were harvested, fixed in formaldehyde and embedded in paraffin. For each animal, 1 every 100 sections (5µm thick, ∼20 per sample) were stained with hematoxylin and eosin (H&E). Metastasis foci and areas were determined by the Histopathology Unit at IGC.

### Statistical analysis

The individual Tumor Control Index (iTCI) was derived from the TCI (16) that compiles three scores assessing tumor inhibition (regression), stability and rejection, per experimental group. The Srivastava Lab. provided a VBA macro with graphical user interface (SL TCI) that automates TCI scores calculation per experimental group. We modified this method to calculate the three sub-scores for individual mice, an improvement that allows for statistical analysis between experimental groups. Statistics of cellular analysis and iTCI were performed using nonparametric Mann-Whitney test. Tumor growth analysis were also performed using two-way ANOVA. Logrank tests were used for survival curves and two-way ANOVA for body weight kinetics. Correlation analyses were performed using Pearson correlation coefficients.

## Acknowledgements

We are grateful to Ana Regalado for production of antibodies, Pedro Faisca’s team for histopathology processing and assessments, Marta Monteiro’s team for maintaining the FACS facility and Manuel Rebelo’s team for mouse husbandry. We thank Thiago Lopes Carvalho and Antonio Coutinho for critical reading of the manuscript. This work was supported by the IGC-Fundação Calouste Gulbenkian and by the Portuguese scientific council (Fundação para a Ciência e a Tecnologia, FCT) including fellowships to JGS (SFRH/BD/52435/2013), IC (SFRH/BPD/111454/2015) and MLB (283/BI/15; UID/Multi/04555/2013). Mouse experiments were in addition supported in part by the national infrastructure CONGENTO (FCT, Lisboa2020, Por2020, ERDF).

## Supplementary Figures legends

**Figure S1.**
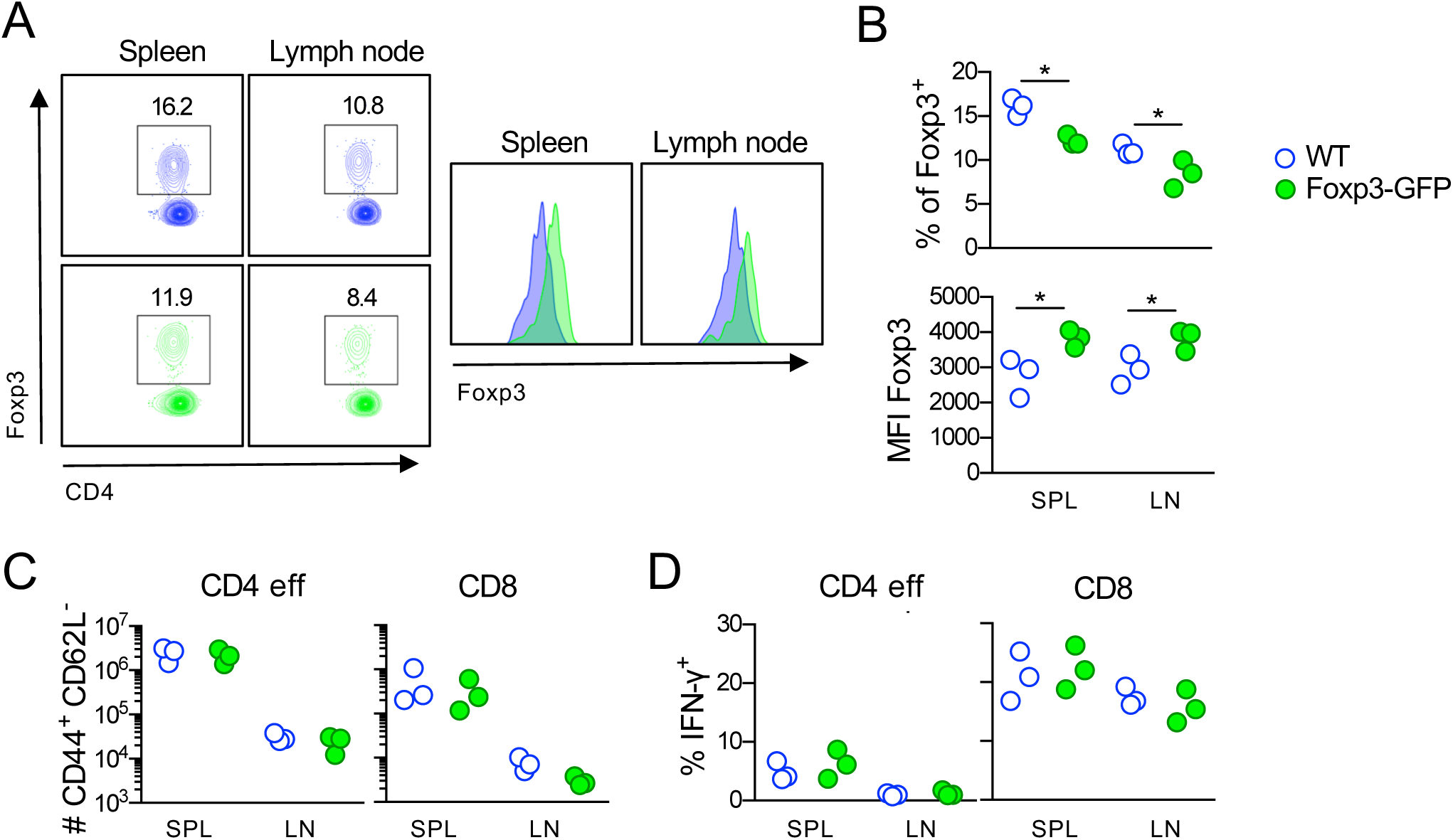
mild cellular alterations at steady state in Ba.Foxp3-GFP mice. Inguinal lymph nodes (LN) and spleen (SPL) from age matched Ba.WT and Ba.Foxp3-GFP mice were analyzed by FACS. **A**) Percentage of i.n. Foxp3^+^ Treg inside gated live CD4^+^ TCR^+^ cells. **B**) Foxp3 mean fluorescent intensity (MFI) in gated Foxp3^+^ cells. **C**) Absolute number of activated (CD44^+^ CD62L^−^) conventional CD4 (Foxp3^−^) and CD8 T cells. **D**) Frequency of Interferon-γ (IFN-γ) producing cells gated on conventional CD4 (Foxp3^−^) and CD8 cells. Statistical analysis performed using nonparametric Mann-Whitney test. *P < 0.05 and **P < 0.005.

**Figure S2.**
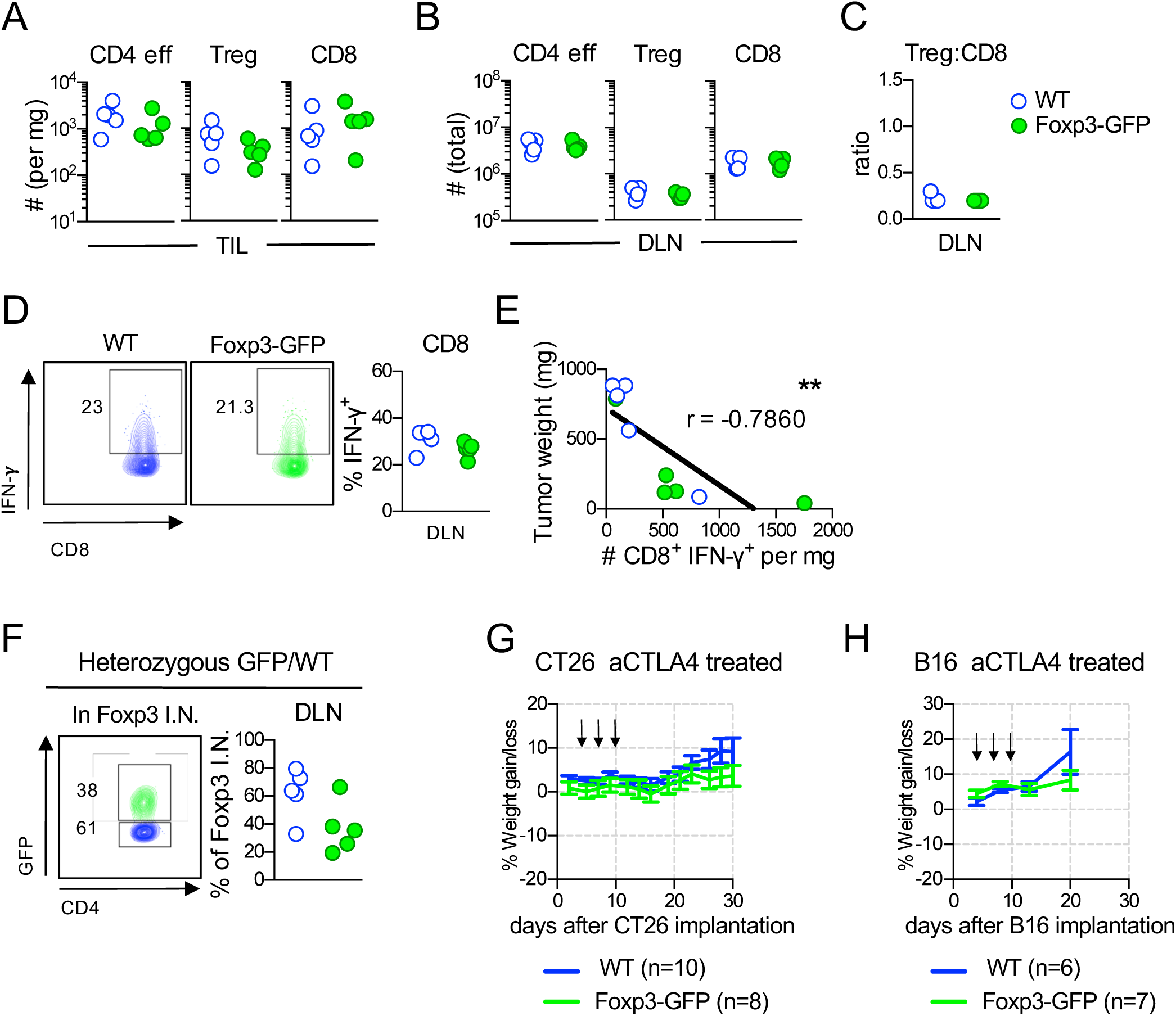
Complement to Figure 1. **A-E**) Complement to cellular analysis shown in Fig1 C-E addressing Ba.WT and Ba.Foxp3-GFP mice implanted with CT26 15 days earlier. A) Numbers of TIL CD4^+^ Foxp3^−^ (CD4 eff), CD4^+^ Foxp3^+^ (Treg) and CD8^+^ (CD8) T cells recovered per mg of tumor. B-C) same mice as in (A) now analyzing CT26 draining lymph nodes (DLN). Sown is absolute number recovered (B), Treg to CD8 ratio (C) and IFN-γ producing CD8 cell (D), from each DLN. E) Correlation between tumor weight and number of TIL IFN-γ producing CD8 T cells per mg of CT26 tumor. **F**) Complement to cellular analysis in Fig. 1F addressing heterozygote Ba.Foxp3^WT/fGFP^ mice. Shown is frequency of GFP expressing cells in gated i.n. Foxp3^+^ cells recovered from DLN. **G, H**) Complement to Fig. 1G and J: Body weight along time in aCTLA4 treated WT and Foxp3-GFP mice implanted with CT26 or B16 tumor. Statistics of cellular analysis were performed using nonparametric Mann-Whitney test. Correlation analysis performed using Pearson correlation coefficients. Body weight kinetic analysis was performed using two-way ANOVA. *P < 0.05, **P < 0.005 and ***P < 0.001.

**Figure S3.**
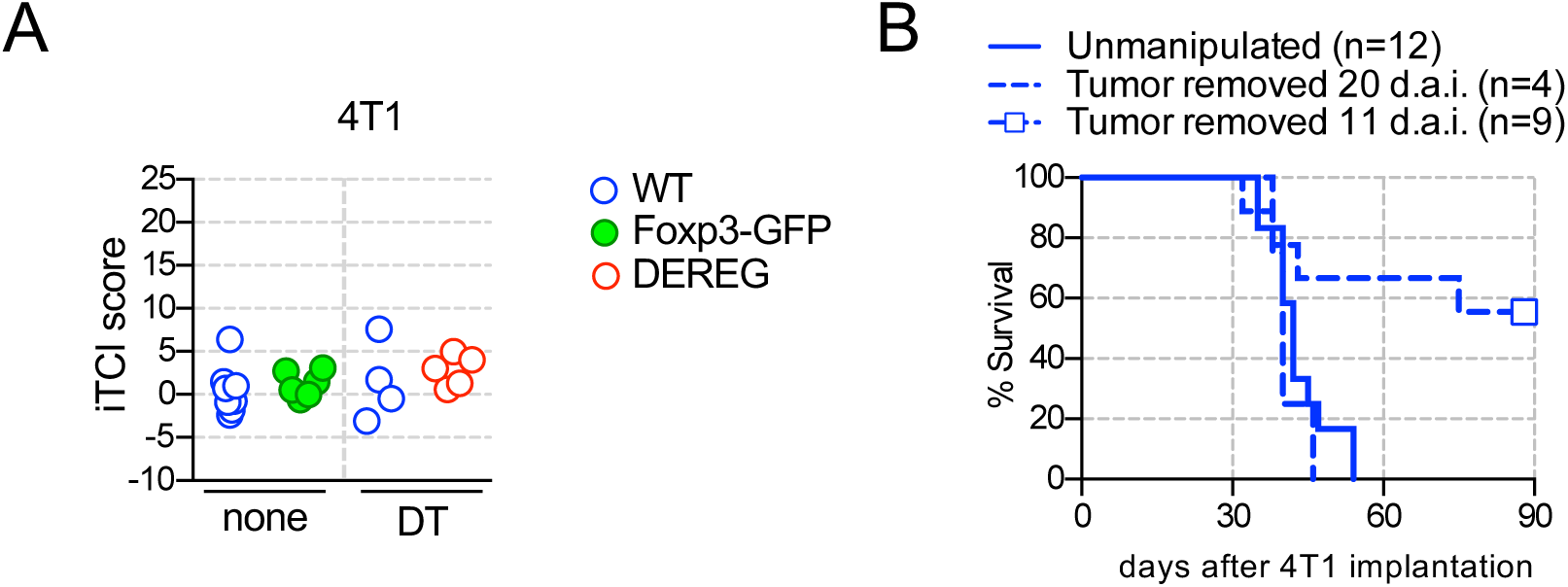
Complement to Figure 2. **A**) iTCI scores (as described in materials and methods) for 4T1 growth curves shown in Fig. 2A and C. Statistical analysis performed using nonparametric Mann-Whitney test. **B**) Survival of WT BALB/c mice submitted to primary tumor resection (doted lines) or not (plain line) at days 11 and 20 after 4T1 implantation (d.a.i.).

